# Temporal clusters of age-related behavioral alterations captured in smartphone touchscreen interactions

**DOI:** 10.1101/2021.12.24.474105

**Authors:** Enea Ceolini, Ruchella Kock, Guido P.H. Band, Gijsbert Stoet, Arko Ghosh

**Affiliations:** Cognitive Psychology Unit, Institute of Psychology, Leiden University, The Netherlands; Department of Psychology, University of Essex, United Kingdom

## Abstract

Cognitive and behavioral abilities alter across the adult life span. Smartphones engage various cognitive functions and the corresponding touchscreen interactions may help resolve if and how the behavior is systematically structured by aging. Here, in a sample spanning the adult lifespan (16 to 86 years, N = 598, accumulating 355 million interactions) we analyzed a range of interaction intervals – from a few milliseconds to a minute. We used probability distributions to cluster the interactions according to their next inter-touch interval dynamics to discover systematic age-related changes at the distinct temporal clusters. There were age-related behavioral losses at the clusters occupying short intervals (~ 100 ms, R2 ~ 0.8) but gains at the long intervals (~ 4 s, R2 ~ 0.4). These correlates were independent of the years of experience on the phone or the choice of fingers used on the screen. We found further evidence for a compartmentalized influence of aging, as individuals simultaneously demonstrated both accelerated and decelerated aging at distant temporal clusters. In contrast to these strong correlations, cognitive tests probing sensorimotor, working memory, and executive processes revealed rather weak age-related decline. Contrary to the common notion of a simple behavioral decline with age based on conventional cognitive tests, we show that real-world behavior does not simply decline and the nature of aging systematically varies according to the underlying temporal dynamics. Of all the imaginable factors determining smartphone interactions in the real world, age-sensitive cognitive and behavioral processes can dominatingly dictate smartphone temporal dynamics.

## Introduction

Quantifying real-world behavior across the adult life span may help unravel the impactful nature of aging and is key to discovering the underlying mechanisms. The gross impact of aging on behavior is widely studied by using self-reports and physical-activity sensors, and they reveal difficulties in the execution of daily behavior and mobility in the elderly (1–4). Real-world behavioral outputs may contain surprisingly informative indicators of aging, such as specific linguistic skills early in life are correlated to healthy cognition later in life (5). Detailed behavioral measurements in the real world may be particularly useful in understanding the interactions between distinct cognitive processes. Indeed, in the naturalistic behavior of typing, the compensatory interplay between hand and eye movements has been observed with aging and this interplay is thought to help maintain typing speed through an otherwise declining motor system (6). Such observations raise the possibility that the impact of aging on real-world behavior may be systematically coordinated – where some types of performance decline while others remain unaffected or even improve, whether this happens under the influence of changing ability or motivation (7–9). However, addressing this possibility in the real world is challenging due to its complex and unclear behavioral structures.

There is emerging evidence that the time series of smartphone touchscreen interactions (tappigraphy) can be used to proxy specific cognitive processes in the real world despite the various ways in which people opt to use their devices – from the distinct body and finger postures to the underlying differences in the motivation for phone use. Tappigraphy contains information on circadian processes, sensorimotor processes, and reward pathways (10–14). The speed of keystrokes on the smartphone declines with age (15). In these recent efforts using tappigraphy, smartphone interaction intervals are simply accumulated and highly reduced before correlating with variables of interest such as age. For instance, the tapping speed may be estimated by using the 25^th^ percentile of the accumulated inter-touch intervals. While this is computationally straightforward, a major limitation is that the intervals are accumulated devoid of contextual information such as on the neighboring intervals. In theory, fast behaviors are driven by fundamentally different cognitive processes than slow behaviors, and a distinct set of processes underlies switching between behaviors (16, 17). Therefore, we introduced the use of joint-interval distribution (JID) to simply structure the time series such that the diverse behaviors are clustered according to the next-interval temporal dynamics (18, 19). For instance, the short inter-touch intervals preceded by similarly short intervals – indicating rapid actions – can be separately considered from the short intervals preceded by long intervals – indicating switching between behaviors with distinct temporal dynamics.

Cognitive tests such as reaction time tests seek to evaluate specific cognitive processes and are routinely deployed to capture age-related decline but they only offer limited insights into real-world behavior. First, the tests may be too detached from how cognitive processes are deployed in real-world behavior (20, 21). For instance, age-related performance decline in the Tower-of-London planning task does not translate to the more ecological Plan-a-Day task (22). Second, many tests – in particular those focused on processing speed – are evaluated in terms of the time taken to perform the task but such emphasis on speed may be present only in a small fraction of real-world behavior. It is notable that in the designed tasks the elderly may be intrinsically inclined to emphasize accuracy at the cost of speed (23). In the context of our analytical framework based on the next intervals on the smartphone, these tasks examine a very narrow temporal segment as in tasks that typically take < 1 s to complete. Here, we seek to address: what are the real-world behavioral correlates of cognitive tests? Do distinct behaviors – occupying the distinct time scales – differently reflect the cognitive tests? Do all fast activities – irrespective of their context – diminish with age? Are age-related real-world behavioral correlates limited to the sub-second time scales?

To address the above questions, we focused on the inter-individual differences in smartphone touchscreen interactions. We clustered the interactions of each individual according to their next-interval temporal dynamics in two-dimensional bins spanning milliseconds to the minute. Towards addressing the implications of the processes captured in cognitive tests for real-world behavior, we correlated the inter-individual differences to cognitive tasks specifically sensitive to sensorimotor processes (visual reaction time), executive functions (task switching), and working memory (Corsi block and 2-back tasks) using mass univariate regressions (24). Notably, the age-related decline of these tasks – albeit at varying degrees – has been previously documented with outstanding declines in the choice reaction time and task switching (25–29). Next, by correlating the interindividual differences in smartphone behavior to chronological age we find clusters of age-related smartphone behavioral correlates such that the correlation strength varies according to the underlying behavioral dynamics. Additionally, the residuals stemming from such regressions can be used to infer accelerated or decelerated aging (30, 31). Using this approach we establish that the same individuals who show accelerated aging in one behavioral cluster can show decelerated aging in another cluster. In our analysis, we considered all smartphone touchscreen interactions and also two subsets of the behavior – when engaged in social networking & browser apps available on the Google play store (henceforth referred to as “Social”) and by focusing on the interactions accumulated when transitioning from one app to the next (henceforth referred to as “Transition”). The latter subsets helped us address if the discovered patterns were also visible in more selective smartphone behavioral data.

## Results

### Distribution of smartphone intervals and the behavioral correlates of cognitive tests

We used joint interval distributions (JIDs) to quantify the probability densities of smartphone behavior at distinct time scales based on all interactions that accumulate within smartphone usage sessions (*Full*, Fig. 1). In addition, we separately considered the interactions on apps used for social networking and the browser (*Social,* Fig. 1). For JID based on intervals involving a switch from one app to the next see supplementary Fig. 1 (indicated as *Transition*). According to grand averages of the ‘*Full*’ and *‘Social’* distributions, the behavior was dominated by short consecutive intervals and by intervals with similar consecutive durations as indicated by the higher probabilities at the corner and the diagonal of the JIDs respectively (the inter-transition intervals were distinctly dominated by long followed by short intervals). Notably, even at the two-dimensional bins with high probabilities, there was substantial inter-individual difference allowing us to relate these variances to the performance in cognitive tests conducted on a PC.

**Figure 1:**
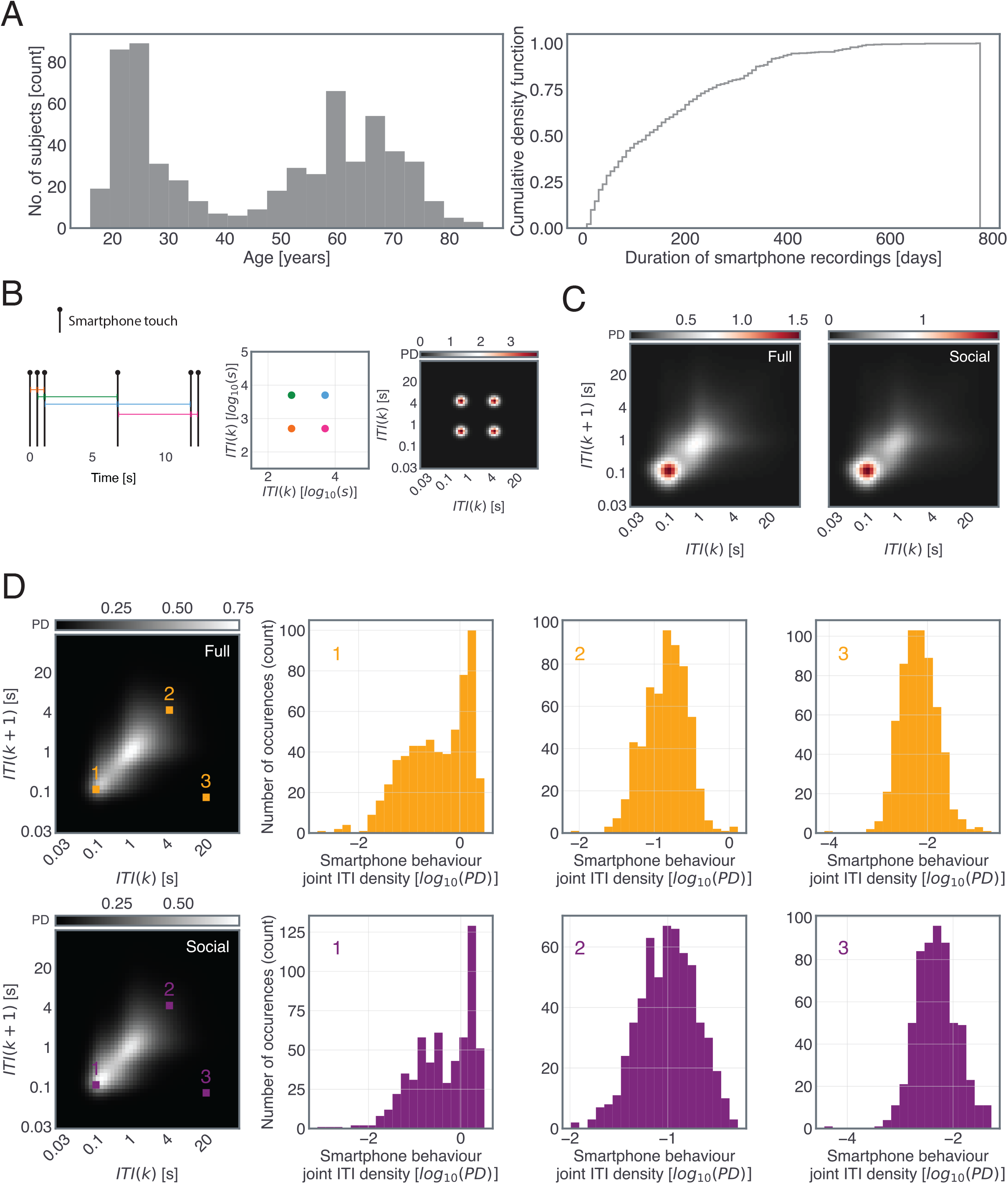
Distribution of inter-touch intervals (ITIs) on the smartphone touchscreen. (**A**) The sample studied here spanned between 16 to 86 years of age. The smartphone recording durations spanned between 7 to 775 days. (**B**) We quantified smartphone behavior using the probability density of joint interval distribution in two-dimensional bins. An example of the probability density resulting from a series of 6 simulated interactions. (**C**) Example joint interval distributions based on all smartphone interactions (Full), and social netoworking and browser apps only (Social) accumulated within smartphone usage sessions spanning 166 days in one individual. (**D**) The population-based mean probability densities and for chosen joint interval bins (yellow and magenta markers) the corresponding inter-individual differences are shown in histograms.

Mass univariate regressions between a cognitive test and all two-dimensional bins of the JIDs were performed to reveal the real-world behavioral correlates of the cognitive tests (Fig. 2, see supplementary Fig. 2 for the age distribution of the participants who were tested on the 2-Back and Corsi Block tests). Towards, the JID smartphone behaviour accumulated within ±10 days of the test were used. For the *Full* JID, the choice reaction time was moderately correlated to a range of smartphone behaviors involving short intervals consecutive to the longer intervals. The slower the reaction time, the lower was the probability densities at the short intervals and the higher were densities at the long intervals. A similar pattern was visible for the *Social* JID (and *Transition* JID, see supplementary Fig. 3). The behavioral correlates for the simple reaction time were weak and mostly constrained in the JID locations occupied by short intervals irrespective of the temporal context, such that the slower the reaction time, the lower the probability densities at the edges of the JID (supplementary Fig. 4). In addition, we found weak correlations for the global cost on the task-switching task for the *Full* and *Social* JIDs (Fig. 2, for Transition JID, see supplementary Fig. 5). No correlations were found for the local cost determined by using the same task. Clusters of behavioral correlates were found for the 2-back tasks such that the higher the sensitivity (*d’),* the higher the probability densities of consecutive short intervals and lower the densities at the longer intervals (supplementary Fig. 6). Stronger correlates were captured for the span on the Corsi Block memory task (supplementary Fig. 7). Overlaying the JID correlation contours of the cognitive tests revealed the differences in the behavioral clusters dominated by the distinct correlates of cognitive tests (Fig. 2).

**Figure 2:**
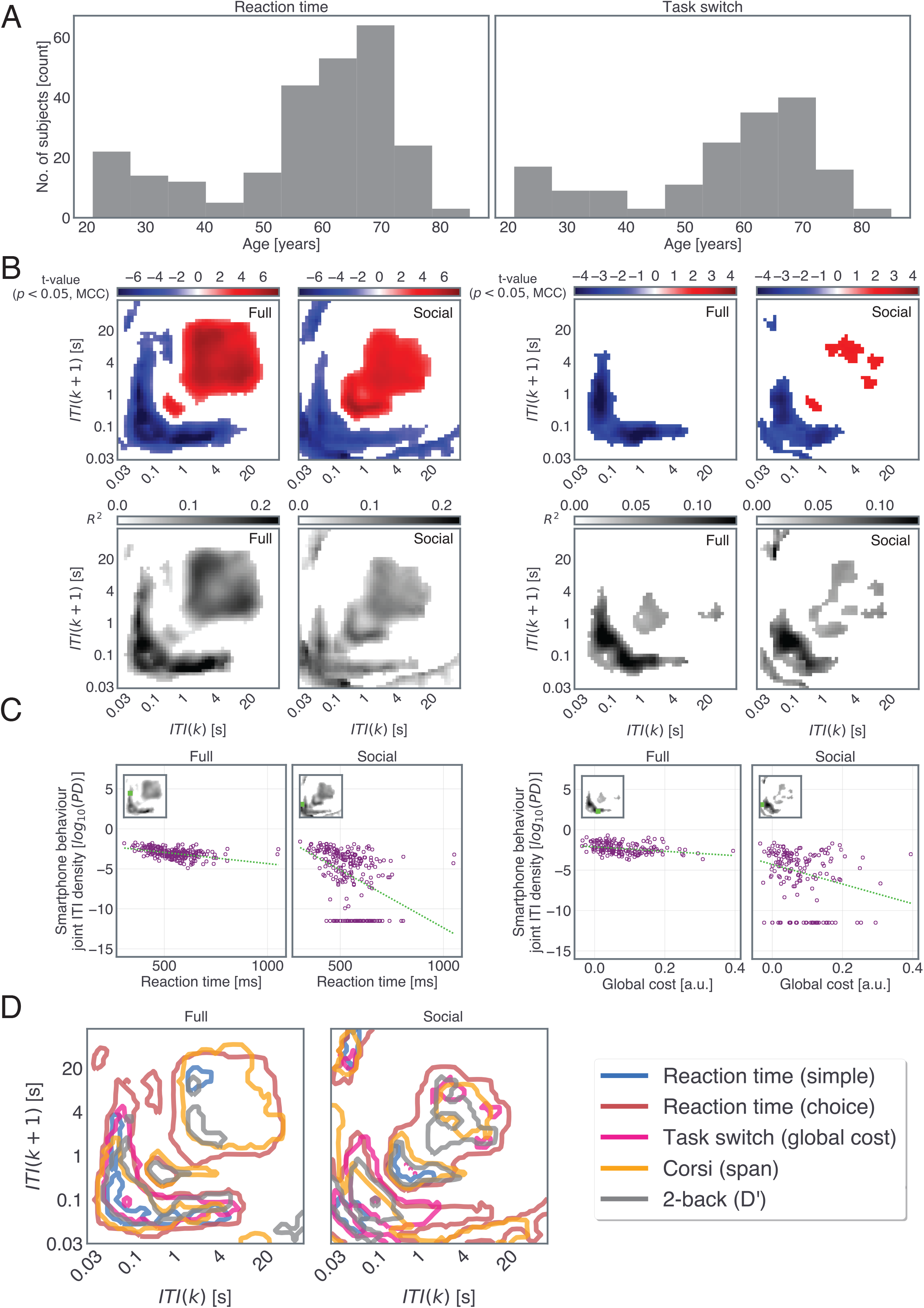
Smartphone behavioral correlates of cognitive tests. (**A**) The distribution of age of the participants who performed the choice reaction time and task switching tests (for the distribution of age corresponding to the other tasks see supplementary Figure 2). (**B**) We performed iterative least-square linear regressions at each two-dimensional bin with cognitive test performance and gender as explanatory variables. The *t*-statistics corresponding to the two tests (left, choice reaction time, and right, global cost estimate on the task switch) are shown in the blue-red color scale when using all smartphone interactions (*Full*) and when using the interactions on social networking apps & browsers (*Social*). The corresponding R^2^ of the full regression model. (**C**) Adjusted response plots to visualize the linear relationships composing the *t*-value representations, with the variables gathered from a single two-dimensional bin. (**D**) Contours of the joint inter-touch interval distributions show significant relationships to the different cognitive tests. The contours lines were thresholded at *f* = 4 for the statistically significant clusters that survived multiple comparison corrections. Note, the taskswitch local costs did not show any significant correlations. All statistics were corrected for multiple comparisons using two-dimensional clustering (1000 bootstraps, α = 0.05).

In sum, all cognitive test outcomes but the local costs on the task-switching test were reflected in the smartphone behavioral distributions. The pattern of the correlations was similar across the distinct forms of JID, but the transition JID showed a weaker spread of correlations. Relative to the other tests, choice reaction time distinctly reflected on a broad range of intervals – spanning from consecutive short intervals to consecutive long intervals and in terms of correlation strength. While multiple tests reflected on the probability densities of consecutive short intervals, only choice reaction time and Corsi Block span consistently dominated the long intervals.

### Age-related smartphone behavioral distributions

We addressed the links between age and the highly reduced measures of smartphone behaviour – usage and entropy based on the JIDs from the entire recording duration; before we describe the correlations against the JIDs’ two dimensional bins. The amount of smartphone usage, quantified as median number of touchscreen interactions decreased with age (*ϐ*_age_ = −0.009; *t*(595) = −14.07; *p* = 4.89 x 10^-39^, *ϐ*_gender (1==male, 2 == female)_ = 0.137; *t*(595) = 5.1200000; *p* = 4.14 x 10^-07^, full model R^2^ = 0.28; *F*(2, 595) = 113.98; *p* = 1.24 x 10^-42^, based on robust multiple regression including gender, supplementary Fig. 8). Additionally, we quantified the behavioral diversity using the entropy of the JIDs (i.e., higher the entropy the more diverse the behaviour). Interestingly, the diversity of the JID marginally declined with age (*Full* JID, *ϐ*_age_ = −0.001; *t*(595) = −2.45; *p* = 0.014, full model R^2^ = 0.08; *F*(2,595); *p* = 7.41 x 10^-12^, based on robust multiple regression including gender, supplementary Fig. 9). Interestingly, this measure showed strong gender differences with males showing more diverse behaviours than females (*ϐ*_gender(1==male, 2 == female)_ = - 0.11; *t*(595) = −5.98; *p* = 3.78 x 10^-9^).

A broad range of the two-dimensional bins of both the *Full* and *Social* JIDs was strongly correlated to age according to the mass univariate analysis linking each JID bin to the chronological age and gender (Fig. 3, for *Transition* JIDs, see supplementary Fig. 10). The probability densities of the short inter-touch intervals were diminished with age whereas longer intervals showed higher probability densities. The correlations were the strongest for the consecutive short intervals and there was a gradual decline in the strength of the correlations as the surrounding intervals became longer (correlation at the bin with max R^2^ on *Full* JID, *ϐ*_age_ = −0.038; *t*(595) = −35.71; *p ≈* 0, full model R^2^ = 0.77; *F*(2, 595) = 639.3; *p ≈* 0, based on robust multiple regression including gender, at a cluster, surviving 2D multiple comparisons correction, scattered in Fig. 3). The *Full* and *Social* JIDs also showed relatively marginal gender differences such that the consecutive short-interval probability densities were higher in females whereas the longer intervals showed higher probability densities in males (supplementary Fig. 11, albeit such patterns, were not present for the *Transition* JID).

**Figure 3:**
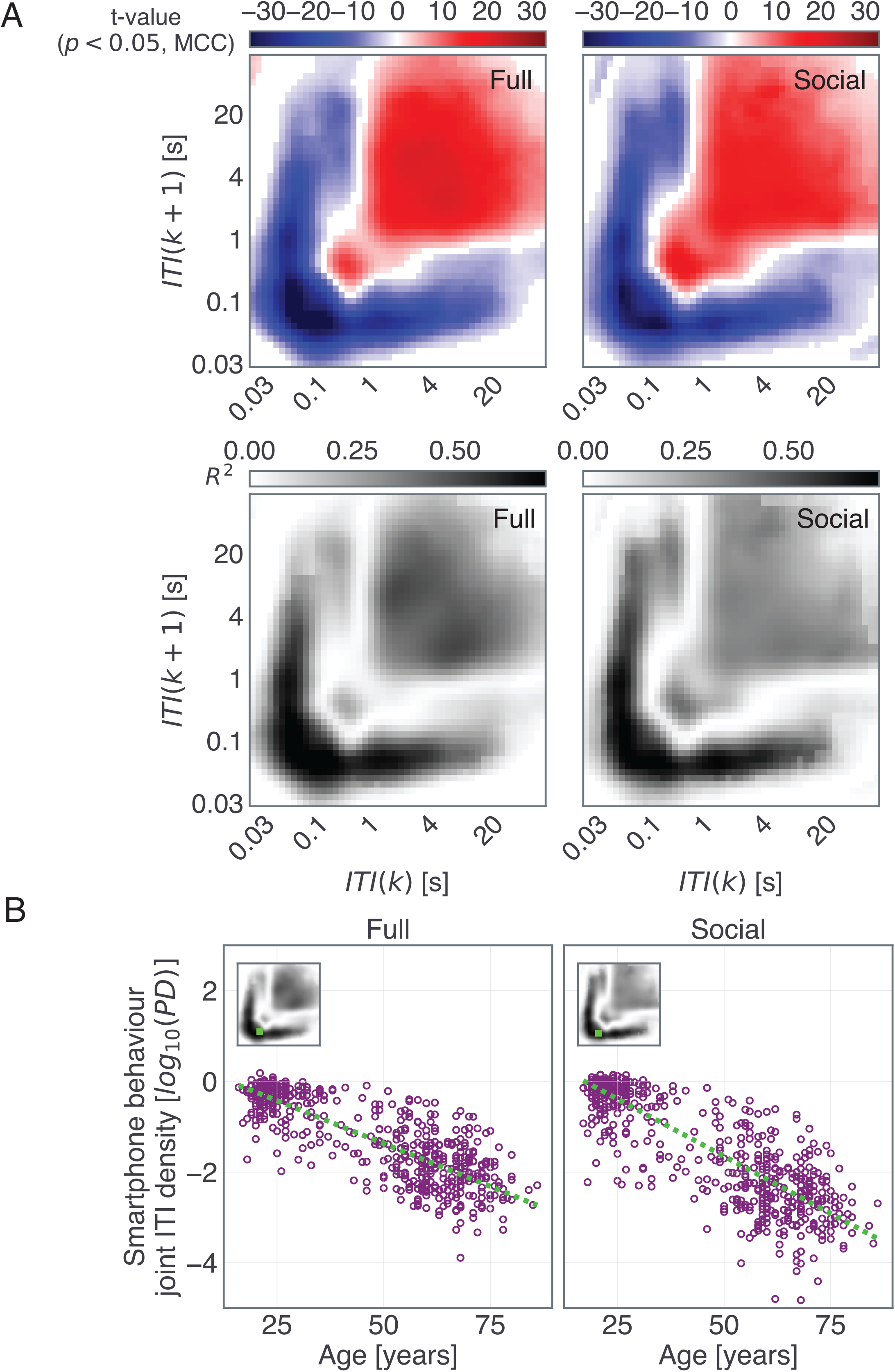
Smartphone behavioral distributions at certain time scales are highly correlated to age. (**A**) We correlated the probability densities at each two-dimensional smartphone behavioral bin with age and gender as explanatory variables. The *t*-statistics (red-blue images) for the variable age reveal substantial behavioral differences across time scales for both types of interval distributions (JID based on all smartphone interactions: ‘Full’ and interactions on social apps and browsers: ‘Social’). The corresponding R^2^ of the full regression model is shown in grayscale images. (**B**) Adjusted response plots of the linear relationships from a select two-dimensional bin (selection noted in the inserted R^2^ plots). All statistics were corrected for multiple comparisons using two-dimensional clustering (1000 bootstraps, α = 0.05). ITI, inter-touch interval, *k*, a given interval, and *k+1* the subsequent interval.

We next addressed whether the pattern of correlations observed in the JIDs was related to the amount of smartphone usage – given that older individuals showed lower amounts of usage. Towards this, we included the amount of usage as a variable in the regression model in addition to age and gender and the overall pattern of age-related correlates remained unchanged due to this inclusion (supplementary Fig. 12). Interestingly, the correlates associated with increased usage were largely confined to the slower intervals. Finally, according to anecdotal observations, older smartphone users may use other digits than the thumb or may be constrained to using only one thumb at a time. We addressed this in a subset of the sample (N = 250) who reported their finger preference on the screen and years of the smartphone experience. Indeed, according to these self reports, the higher the age, the lower the reliance on the thumb (*ϐ*_Thumb pref_. = −0.0012, *t*(247) = −3.55; *p* = 0.00046, full model R^2^ = 0.087; *F*(2, 247) = 11.7; *p* = 1.38 x 10^-5^, based on robust multiple regression including gender) and the lower the simultaneous use of both thumbs (*ϐ*_Dual thumb use_ = - 0.0032, *t*(247) = −3.75; *p* = 0.00022, full model R^2^ = 0.063; *F*(2, 247) = 8.2; *p* = 0.00035, based on robust multiple regression including gender). Including these finger usage variables and the years of smartphone experience in the regression model linking smartphone JID and age, did not alter the pattern of age-related behavioral correlates (supplementary Fig. 13).

In sum, chronological age explained a substantial extent of the inter-individual differences in smartphone behavior captured using the JIDs. The probability of short consecutive intervals was particularly depleted with advanced age whereas the probability of longer intervals was higher with increased age. These correlates were independent of how much people used their devices, the number of years spent on the smartphone, and the choice of the fingers used on the touchscreen.

### Chronological age is reflected in cognitive tests but weaker than in smartphone behavior

We next addressed the extent to which chronological age reflected in cognitive tests (supplementary fig. 14). The simple reaction time was only weakly correlated with age (*ϐ*_age_ = 0.65; *F*(2,256) = 8.77; *p* = 0.0033, R^2^ = 0.034, Robust simple regression corrected for gender). The choice reaction time showed a stronger relationship than simple reaction time, albeit still moderate in terms of correlational strength, such that the higher the age the slower the reaction time (*ϐ*_age_ = 3.42; *F*(2,257) = 101.54; *p* ≈ 0; R^2^ = 0.28, Robust simple regression corrected for gender). The global cost (but not local cost) on the task switching task was marginally correlated with age (*ϐ*_age_ = 0.001; *F*(2,166) = 10.19; *p* = 0.0017; R^2^ = 0.058, Robust simple regression corrected for gender). The corsi span too declined with age (*ϐ*_age_ = −0.0305; *F*(2,239) = 54.56; *p* = 5.21 x 10^-12^; R^2^ = 0.18, Robust simple regression corrected for gender). There was a tendency of age-related decline for the 2-back D’ (*ϐ*_age_ = - 0.018; *F*(2,189) = 2.94; *p* = 0.09; R^2^ = 0.017, Robust simple regression corrected for gender).

The best performing cognitive test in terms of capturing age-related decline was the choice reaction time (R^2^ = 0.28). Next, we contrasted this regression to the age-related decline captured on JID. Towards this, we focused on the subset of the participants who performed the choice reaction time task and re-estimated the smartphone correlates of age (adjusted for gender). In this subset, the best related two-dimensional bin (in terms of R^2^ on the *Full* JID) was superior to that of the regression capturing choice reaction time (*ϐ*_age_ = - 0.046; *F*(2,257) = 353.57; *p* ≈ 0; R^2^ = 0.58, Robust simple regression corrected for gender, at a cluster surviving 2D multiple comparisons correction).

### The distinct pace of aging in the different behaviors captured on the smartphone

The nature of the age-related correlations varied from one type of smartphone temporal dynamics to another. The resulting linear models were further leveraged to gain insights into the pace of aging. Considering that the regression fitted age vs. probability density at a two-dimensional bin as an indication of typical aging, we used the deviations from this fit (residuals) to quantify accelerated or decelerated aging (for *Full* and *Social* JIDs see Fig. 4, for Transition JID see supplementary Fig. 15). To elaborate, at a temporal bin showing an age-related reduction in probability densities, positive residuals indicate decelerated aging whereas negative residuals indicate accelerated aging. We performed mass cross-correlations between the age-related residuals between pairs of the two-dimensional bins of the JIDs to identify two forms of relationships: consistent age-related relationships, as in where accelerated aging in one two-dimensional bin also indicated accelerated aging in another bin, and inconsistent relationships, as in where accelerated aging in one bin indicated decelerated aging in another bin. For both *Full* and *Social* JIDs, consistent relationships dominated the symmetrically opposite (say short followed by long intervals and long followed by short intervals) two-dimensional bins. However, the inconsistent relationships revealed links between behaviors with rather distinct next interval temporal dynamics (such as along the diagonal of the JID).

**Figure 4:**
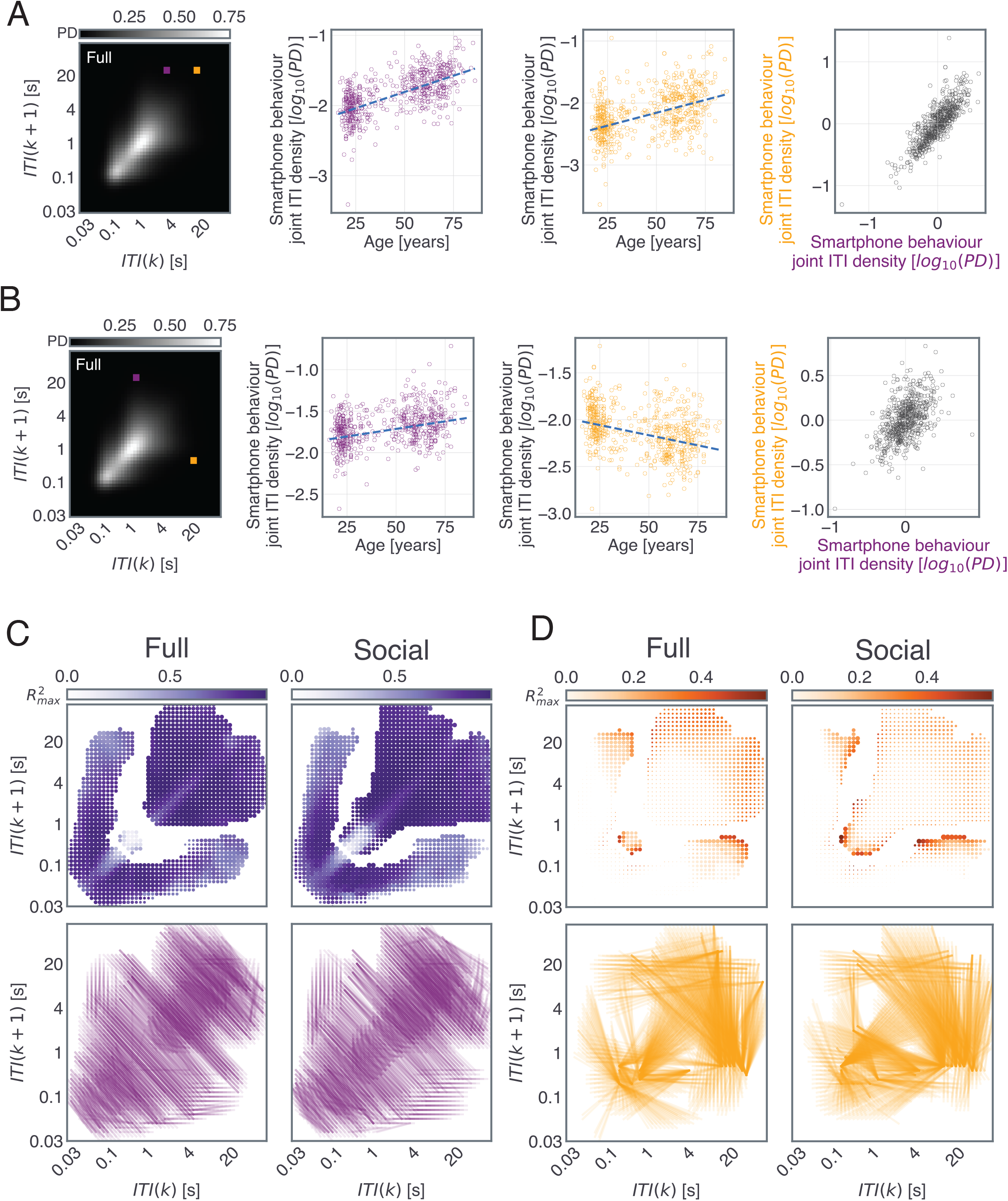
Locating consistency of aging across different temporal segments of smartphone behavior. We used pair-wise regressions relating the age-residuals across smartphone behavioral bins. (**A**) In this example pair, the age-residuals of the two-dimensional bins are correlated to each other revealing a consistency – so individuals who show accelerated aging in the first two-dimensional bin (magenta) also show accelerated aging in the second bin (yellow). (**B**) In this example pair, the age-residuals of the two marked bins are correlated to each other revealing an inconsistency. Here, in the first two-dimensional bin (magenta) individuals who had lower behavioral probability density than the fitted line were considered as better performers (as this behavioral density increases with age, positive slope). In the second two-dimensional bin (yellow) individuals who had higher behavioral probability density than the fitted line are considered better performers (as this behavioral density reduces with age, negative slope). The regression between the residuals shows a positive correlation – so individuals with accelerated aging in one bin show decelerated aging in another. (**C**) All two-dimensional bins with significant age-related correlations were selected for the residual analysis. Here we show the max R^2^ for the *consistent* pairs observed at a given twodimensional bin. The corresponding line plots (each line depicting a significant correlation) show the location of the *consistent* correlational pattern linking the residuals, where the bins were separated by a bin distance of > 5 bins. (**D**) Same as ‘C’ but for *inconsistent* pairs.

## Discussion

Smartphone interactions reflected chronological age in a sample of smartphone users spanning the adult lifespan. The direction and strength of the age-related correlations varied according to the behavioral dynamics. While the consecutive short inter-touch intervals were diminished with age, the longer intervals were enhanced. The putatively distinct impact of aging across smartphone behavior may be attributed to the status of the underlying cognitive functions and shifting behavioral strategies. Our follow-up analysis suggests a form of compartmentalization such that distinct smartphone behaviors age in different directions – to deplete or to raise the probability density – within the same individual.

We addressed the possibility that the age-related correlations in smartphone behavior were shaped by commonly considered or intuitive factors such as the amount of phone usage, years of experience, and the choice of fingers used on the phone. The amount of smartphone usage and fingers used were weakly related to the chronological age confirming anecdotal observations that older individuals may use their phones less and differently. Nevertheless, including these factors in the multivariate regression model along with chronological age did not alter the pattern of age-related correlations in the smartphone behavioral JIDs. This suggests that the age-related correlates discovered here may be attributed to other factors beyond experience and postural choices. The limitation of experience and finger choices was also evident in the form of near absent correlations of these factors across the smartphone interactions captured in the joint interval distribution.

The weak or marginal correlations linking chronological age and overall smartphone usage is in line with prior observations (32). In contrast to the links to usage, we found strong correlations when considering the probability densities clustered according to the next interval dynamics. This underscores the importance of capturing the temporal dynamics underlying real-world behavior for aging research. Moreover, the overall pattern of relations was visible in the JIDs using only a subset of the data – that is when using the data from social networking & browser apps (*Social* JID), or when considering only those interactions made upon transitioning to an app (*Transition* JID). Notably, the shortest intervals in the *Transition* JID were in the range of ~1 s whereas for the *Full* and *Social* JIDs the shortest intervals were in the range of ~100 ms. Despite this temporal shift the overall pattern of age-related correlates remained consistent such that the short intervals in the respective JIDs declined with age whereas the longer intervals were enhanced. This suggests that the relative instead of the absolute temporal dynamics have a putative role in how the behavior is shaped by aging.

Intriguingly, both the *Full* and *Social* distributions were dominated by behaviors involving similar consecutive intervals across time scales (to occupy regions around the diagonal of the JID). This pattern may be an outcome of behavioral states reliant on similar consecutive outputs such as in reading a webpage by using consecutive swipes of similar intervals. Under the assumption that shorter intervals reflect texting or rapid browsing without absorbing information, and longer intervals reflect browsing or reading, perhaps with age individuals engage on the smartphone to absorb more information rather than generate rapid outputs. The idea that there may be overall changes in behavioral structures or strategies is further supported by the age-related decline in the JID entropy. This loss of behavioral complexity may parallel the age-related loss of physiological complexity as captured using time-series measures such as in fluctuations of the heart rate (33, 34).

The regressions between the cognitive tests and smartphone behavior provide insights into how the intrinsic cognitive properties may shape daily behavior. Simple reaction time has been associated with measures from daily life such as falls in the elderly and questionnaires that probe daily action outcomes (35, 36). In our analysis, we found that simple reaction time was mostly reflected in the probability of behavior with short inter-touch touch intervals – irrespective of their neighboring temporal context. This suggests that the cognitive processes reflected in simple reaction time specifically impact those daily behavioral outputs that depend on rapid cognitive processing. This was in contrast to the smartphone behavioral correlates of choice reaction time which included both the short and long intervals. These distinct patterns provide real-world behavioral support to the long-standing idea that cognitive processes underlying simple reaction time are more constrained than the processes underlying choice reaction times (37). The behavioral correlates of the Corsi span too broadly occupied short and long intervals, and the correlates were more widespread than for the 2-back D’. This indicates that visual-spatial memory has a more prominent role in smartphone behavior than verbal memory. There was also a sharp contrast between the pattern of results for local vs. global switch costs. Local costs were unrelated to smartphone behavior whereas global costs were linked to the short intervals (and long intervals on the *Transition* JID). The later findings based on global costs underscore the real world relevance of maintaining multiple task configurations. The former findings raises the possibility that processes captured by the local switch costs may be task specific and do not play a dominant role in real world behaviour. The two-dimensional bins at the short consecutive intervals were commonly correlated to these diverse tests. Perhaps multiple cognitive processes culminate to enable the generation of sustained rapid actions in daily life.

According to our analysis, the short consecutive inter-touch intervals were particularly vulnerable to aging. The variance in age explained by this segment of smartphone behavior was superior to the variance explained by any of the cognitive tests aimed at specific cognitive processes. We speculate that the putative reliance on multiple cognitive processes to generate actions with short consecutive intervals also makes the same behavioral outputs more vulnerable to age than other actions. According to previous reports, elderly typists engage compensatory strategies – such as faster eye movements – to protect typing speed against age-related decline (6). Our data suggest that such compensatory strategies are limited.

We considered the residuals stemming from the smartphone JID correlations with age as a behavioral measure of accelerated or decelerated aging. Accelerated aging at certain temporal scales of the JID was correlated to decelerated aging at others and such inconsistent links were present across the time scales. Essentially, the same individuals could show divergent forms of aging at the different temporal bins. This helps further extend the idea that the nature of aging varies according to the underlying temporal dynamics. Taken together with the correlations of cognitive tests suggesting that distinct temporal locations of the JID are driven by distinct cognitive processes our findings are in line with the emerging evidence showing different forms of aging in the distinct cognitive processes (8).

The pattern of results solely based on the cognitive tests mimicked previous reports on the age-related decline. As observed before, the simple reaction time was less sensitive to aging in comparison to the choice reaction time (25, 29, 38). Similarly, the local cost on the task-switching test was less sensitive to aging than the global cost (27). Here, as in previous reports, chronological age and gender explained only a small part of the inter-individual variance in the cognitive tests (39, 40). While it is well established that cognitive test performance contains information on the aging status, our results indicate that specific aspects of smartphone behavioral output may contain even stronger indicators of age. Real-world behavioral data may be as informative (if not more) on aging as the cognitive tests.

Our study demonstrates that smartphone interactions offer unique insights into how daily life is shaped by aging across the adult life span. Our study helps upgrade the accounts of age-related decline based on narrow and artificial tasks with accounts of systematic age-related behavioral organization in daily life occupying a broad range of behavioral dynamics captured on the smartphone. These findings pave the way for longitudinal studies to track the status of aging at the level of each individual and across a broad range of behavioral dynamics. Inter-individual differences in real-world smartphone behavior – that is intuitively under diverse, long, and difficult to quantify influences such as body posture, prior experience, goals, decision making, impulsivity, habits, and information content on the screen, to list a few – maybe instead dominated by hidden cognitive and behavioral processes under the influence of aging.

## Methods

### Participants

We recruited 720 volunteers older than 16 years of age through on-campus flyers and emails, and via the agestudy.nl data collection platform aimed at the general population in The Netherlands. The recruitment to the online platform was facilitated by hersenonderzoek.nl which contains a list of willing research participants in The Netherlands (41). The data from volunteers based on the on-campus recruitment have been used in previous reports (11, 14). Only self-reported healthy participants in terms of neurological and psychological health (at the time of recruitment) were considered. Additional exclusion criteria included individuals who shared their smartphones with others, had dysfunctional or missing digits of the hand, people without access to email, and a desktop or laptop (for cognitive tests). All participants gave informed consent either via paper signatures or by using a digital form. The experimental procedures were approved by The Psychology Research Ethics Committee at Leiden University.

Of the 720 recruits, 642 successfully installed the smartphone data logging app and reported their age in terms of the month and year of birth (235 males, 396 females, 11 unreported or unclear gender reports, 16 to 86 years of age). From this cohort, we analyzed 598 participants based on a cut-off of 7 days of smartphone recordings and a minimum of 100 smartphone interactions in that period.

### Online data collection platform

We constructed a cloud-based (IBM Cloud, IBM, Armonk) data collection platform (agestudy.nl) to anonymously link the data – self-reported age and gender, cognitive tests, and smartphone data. They were linked through unique personal identification numbers to each participant upon sign-up. The platform presented the study information, enabled users to sign-up for the study using an online form, and recorded the age and gender self-reports, smartphone usage, and finger use questionnaire, in addition to other questionnaires as a part of larger data collection efforts (not considered here). The sign-up process was supported via email and phone consultations on demand. The platform enabled users to access the cognitive tests implemented on psytoolkit.org (42, 43).

### Finger configuration preference estimation

Participants reported on their smartphone behavior using a questionnaire. The question of interest was: ‘How many years have you used a smartphone?’. The users ranked their finger preference by sorting 8 images from most used to least used. The images represented: left thumb, right thumb, left&right thumb in portrait mode, left&right thumb in landscape mode, left index finger, right index finger, left middle finger, and right middle finger. These ranks were converted into two scales ranging from 0 to 1. With the first scale capturing the preference for the thumb, with 1 indicating the thumb was most preferred, and the second scale capturing the preference for simultaneous use of both thumbs, with 1 indicating dual use was most preferred.

### Cognitive tests

Participants performed the cognitive tests online via agestudy.nl and psytoolkit.org, and these tests were activated when seated in front of a laptop or personal computer with a keyboard (self-reported). The tests were administered in English or Dutch. When logged in participants could choose between Deary-Liewald reaction time tasks, cued task-switching, Corsi block tapping, and 2-back tasks, apart from questionnaire-based tasks. The participants were instructed to perform at least one task per month.

The Deary-Liewald reaction time tasks were used to measure simple and choice visual reaction times (38). Towards simple reaction time, participants were instructed to respond to the presence of a checked square using a keypress (key K) with the right index finger as fast as possible. An instruction video and a practice block of 8 trials proceeded with the main task consisting of 25 trials. In the choice reaction task, we used a 4-choice paradigm. The participants were instructed to respond to the presence of a checked mark (in one of the 4 squares) using the middle or index fingers of the left or right hands corresponding to the location (keys S, D, K, and L). A practice block of 8 trials proceeded with the main task consisting of 50 trials. In both tasks, participants were informed if they were too slow (> 3 s) or if they used incorrect keys on a given trial. The inter-trial interval was set within the range of 1 s and 3 s.

We used the Corsi Block tapping task to evaluate short-term spatial memory (44). Briefly, nine spatially distributed locations on the screen were highlighted in certain sequential patterns and the participants were instructed to report the presented sequence by recreating the same order. The number of locations was progressively increased to determine the Corsi span until two consecutive errors were made or when all the 9 positions were correctly ordered. Participants were presented with an instruction video and performed a practice set before the main test.

We used a 2-back task to evaluate working memory. A train of letters was presented and the participant responded with a keypress (key M) when the same letter was shown two trials back. The practice set was customized such that the previously shown letters were still visible to the user in a lighter shade. A video tutorial was followed by a practice set containing 15 trials leading to the main task in three blocks of 30 trials. The gap between stimuli presentation was set at 2 s and each stimulus was presented for 760 ms. Correct responses triggered visual feedback.

A cued task-switching shape-color task was used. In the shape task, participants distinguished between circles and squares, and in the color task, participants distinguished between yellow and blue (using the keys D and K). A measurement session consisted of (in order) an optional video tutorial, a block of color tasks, a shape task, two mixed blocks, followed by a shape and color block again. The first blocks of each type were preceded by a set of 8 training trials. The color or shape blocks consisted of 20 trials each, and the mixed blocks consisted of 50 trials. The type of the task was indicated with a text cue as in ‘Shape’ vs. ‘Color’. The cue was displayed for 400 ms and cue to item delay was set at 500 ms. Subsequent trials were triggered after waiting for 5 s for a response.

For inclusion in the data analysis a minimum number of trials had to be successfully executed per task: 12 trials, simple reaction time; 24 trials, choice reaction time; 12 trials, task switching task; two trials, Corsi; 30 trials, 2-back. Individuals who did not fulfill one or more tasks were retained for the analysis involving the other tasks. Median reaction times were used to quantify the simple reaction time, choice reaction time, global and local cost. The global cost was normalized using the median reaction time of the shape and color blocks. The local cost was normalized using the median reaction time of the non-switch trials. Corsi span was used to quantify the performance on the Corsi block tapping task. Sensitivity (D’) was used to quantify the 2-back performance.

### Smartphone data collection

We use a background App and a cloud-based data collection service to record the smartphone touchscreen interactions (TapCounter, QuantActions Ltd, Lausanne). Briefly, the app was downloaded and installed from the Google Playstore (Google, Mountain View). The participant entered a unique ID into the App to link with the other data. Through the data collection period, the incoming data was monitored via a web-based management tool (Taps.ai, QuantActions Ltd, Lausanne). The raw smartphone data was downloaded along with the unique ID and parsed using MATLAB (Mathworks, Natick). The parsed data included the timestamp of the touchscreen interactions, the label of the foreground app in use, the timestamp of the screen on and off events. We quantified the amount of usage (log10 count) as the median number of touchscreen interactions accumulated per day, with the days where no touches were recorded excluded from the median.

### The joint interval distribution (JID)

The inter-touch intervals were quantified in terms of joint-interval distributions to quantify the next interval dynamics of smartphone interactions as introduced before (18). Only those intervals gathered with the screen on were considered, i.e., intervals between one usage session (period from the screen turning on to being switched off) to the next were not considered here. The JID was based on subsequent inter-touch intervals (ITI). Any ITI (say *k*) was related against the subsequent interval (say *k* + 1). By using kernel density estimation over the accumulated ITIs we estimated the joint probabilities on *log10* transformed data. A Gaussian kernel with a bandwidth of 0.1 was used towards the two-dimensional probability density estimation (45). The output of the kernel density estimation was then discretized using 50 bins per dimension in the range of 10^0.5^ ms and 10^5^ ms. The range was determined based on the 99^th^ percentile of all the ITIs across all the subjects. The entropy of JID was estimated using the information theory entropy formula in 2D.

We estimated three different forms of JID. In the first form, we used all accumulated intervals (‘Full’). In the second form (‘Social’), we only used those intervals accumulated while on social networking and browser apps as defined according to the Google play (Google, Mountainview) ‘Communication’ category (such as WhatsApp, Facebook, Chome, available through Taps.ai, QuantActions, Lausanne). In the third form (‘Transition’) we considered the intervals based on the interactions accumulated while transitioning from one app to another. For the regression models linking the JIDs with age and gender, the data from the entire recording period was used. For the regression models linking JIDs with the cognitive test, the smartphone data from the ± 10 days from the day of the test was used, and subjects who did not have data in this period were eliminated.

### Mass univariate regression analysis

We used the hierarchical linear modeling toolbox LIMO EEG for regression analyses and multiple comparison corrections (24). To link the JID to cognitive tests we performed robust linear regressions (iterative least squares) linking each two-dimensional JID bin to the test output and gender (dummy variable). The probability densities were log10 transformed, with the minimum value encountered in the population replacing any zero. The resulting statistics were corrected for multiple comparisons using 2D clustering implemented in LIMO EEG (*α* = 0.05, 1000 bootstraps). The same approach was used when linking to age, and additional models were raised with usage as a variable.

To study the age-related residuals, we first isolated the residuals in a two-step manner with the first step involving a linear regression using the gender dummy variables, and a subsequent linear regression against age. In the study of consistency of age-related residuals within the JID we performed pairwise correlations across all bins at a distance greater than 5 bins, R^2^ > 0.1, and survived 2D multiple comparison correction, in the second regression. The resulting analysis was corrected for multiple comparisons using the false discovery rate implemented in LIMO EEG (*α* = 0.001).

### Code availability

The codes used to analyze the links between JID, age, and cognitive tests are (upon publication) deposited on the CODELAB git repository. Furthermore, the codes used for cognitive tests are available on request without reason from the corresponding author via psytoolkit.org.

### Data availability

The analyzed data from the level of the JID and the associated cognitive test results, age, and gender information is made available on dataverse.nl (unlocked upon publication).

## Acknowledgments

We are grateful to Eva van Workum and Irene Taboada Warmerdam for their communication efforts with the participants. We thank Sven van Es, Charlotte van der Berg, Bo Bekkers, and Michelle Snoeks for helping construct the agestudy.nl tasks and instructions. We thank Richard Ridderinkhof for the suggested inclusion of finger preference parameters in the analysis. We thank Wen Yu Wan for coding the script to extract information on fingers used on the phone. We thank all the participants who contribute to this study with their time.

## Figure legends and associated supplementary figures

Supplementary Figure 1. *The Transition JID.* Population means of the probability densities derived at each two-dimensional bin.

Supplementary Figure 2. *The distribution of age of the participants who performed the Corsi block and 2-back tasks.*

Supplementary Figure 3. *Transition JID correlates with the choice reaction time.* The t-statistics correspond to the choice reaction time (red-blue image) and the corresponding R^2^ of the full regression model (incl. gender, gray image). The statistics were corrected for multiple comparisons using two-dimensional clustering.

Supplementary Figure 4. *Smartphone correlates of simple reaction time.* The smartphone behavioral correlates of simple reaction time on the Full, Social, and Transition JIDs. The t-statistics correspond to the cognitive test (red-blue image) and the corresponding R^2^ of the full regression model (incl. gender, gray image). The statistics were corrected for multiple comparisons using two-dimensional clustering.

Supplementary Figure 5. *Transition JID correlates of global cost.* The smartphone behavioral correlates of the global cost on the Transition JID. The t-statistics correspond to the cognitive test (red-blue image) and the corresponding R2 of the full regression model (incl. gender, gray image). The statistics were corrected for multiple comparisons using two-dimensional clustering.

Supplementary Figure 6. *Smartphone behavioral correlates of the 2-back test.* The smartphone behavioral correlates of 2-back D’ on the Full, Social, and Transition JID. The t-statistics correspond to the cognitive test (red-blue image) and the corresponding R^2^ of the full regression model (incl. gender, gray image). The statistics were corrected for multiple comparisons using two-dimensional clustering.

Supplementary Figure 7. *Smartphone correlates of Corsi Block.* The smartphone behavioral correlates of the span on the Corsi Block task on the Full, Social, and Transition JID. The t-statistics correspond to the cognitive test (red-blue image) and the corresponding R^2^ of the full regression model (incl. gender, gray image). The statistics were corrected for multiple comparisons using twodimensional clustering.

Supplementary Figure 8. Smartphone correlates of age. The amount of smartphone usage correlates with age. Adjusted response plot showing the linear relation between smartphone usage vs. age. Based on robust linear regression against age and gender.

Supplementary Figure 9. Smartphone entropy. The entropy of the JID is related to age. The Full, Social, and Transition JIDs with min and max entropy values in the sampled population. The adjusted response plots show the linear relation between the corresponding entropy vs. age. Based on robust linear regression against age and gender.

Supplementary Figure 10. Transition JID correlates of age. The correlation between Transition JID and age. Legend same as in Fig. 3 but for Transition JID.

Supplementary Figure 11. Smartphone correlates of gender on the JIDs. The t-statistics (red-blue images) for the variable gender (dummy variable in the regression model including age). Male 1, and female 2 were used as dummy variables. Note, females (2) showed higher probability densities of short consecutive intervals.

Supplementary Figure 12. *A multivariate model including age, gender, and smartphone usage linking to the JIDs.* The results of mass univariate regression include the amount of smartphone usage in terms of the median number of interactions per day in addition to age and gender in the mass regressions conducted at each two-dimensional bin. The t-statistics for all of the variables and corresponding R^2^ of the full model. All statistics were corrected for multiple comparisons using twodimensional clustering (α = 0.05).

Supplementary Figure 13*. Age-related correlates in JIDs remain in the presence of other smartphone use-related variables.* We used a multivariate model to explain the inter-individual differences in the JID and this model included age, gender, smartphone usage, thumb preference score, dual thumb preference score, and the years of the smartphone experience. The t-statistics for all of the variables and corresponding R^2^ of the full model. All statistics were corrected for multiple comparisons using two-dimensional clustering (α = 0.05).

Supplementary Figure 14. *Age-related correlates of cognitive tests.* The adjusted response plots are based on multivariate regression models containing the variables age and gender. Regression fits with *p* < 0.05 for the variable age are plotted.

Supplementary Figure 15. *Accelerated & decelerated aging captured on Transition JID.* (**A**) Consistent relationships and the corresponding links and (B) inconsistent relationships and the corresponding links. Legend same as in Figure 4 C & D.

