## Supplementary Figures for "Temporal clusters of age-related behavioral alterations captured in smartphone touchscreen interactions"

### Smartphone interactions reveal the loci of age-related behavioral changes

December 3, 2021

#### List of Supplementary Figures

1 The Transition JID. . . . . . . . . . . . . . . . . . . . . . . . . . .
2 The distribution of age of the participants who performed the Corsi block and 2-back tasks. . . . . . . . . . . . . . . . . . . . .
3 Transition JID correlates with the choice reaction time. . . . . .
4 Smartphone correlates of simple reaction time. . . . . . . . . . .
5 Transition JID correlates of global cost. . . . . . . . . . . . . . .
6 Smartphone behavioral correlates of the 2-back test. . . . . . . .
7 Smartphone correlates of Corsi Block. . . . . . . . . . . . . . . .
8 Smartphone correlates of age. . . . . . . . . . . . . . . . . . . . .
9 Smartphone entropy. . . . . . . . . . . . . . . . . . . . . . . . . .
10 Transition JID correlates of age. . . . . . . . . . . . . . . . . . .
11 Smartphone correlates of gender on the JIDs. . . . . . . . . . . .
12 A multivariate model including age, gender, and smartphone usage linking to the JIDs. . . . . . . . . . . . . . . . . . . . . . .
13 Age-related correlates in JIDs remain in the presence of other smartphone use-related variables. . . . . . . . . . . . . . . . . . .
14 Age-related correlates of cognitive tests. . . . . . . . . . . . . . .
15 Accelerated & decelerated aging captured on Transition JID. . .

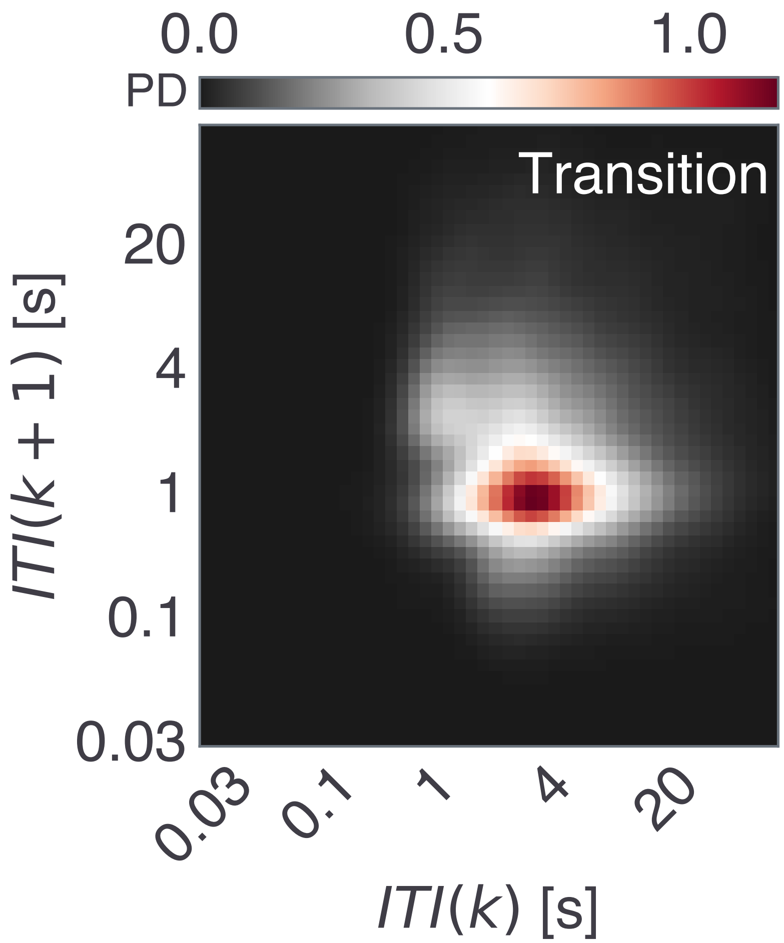


Supplementary Figure 1: The Transition JID.

Supplementary Figure 1: Population means of the probability densities derived at each two dimensional bin.

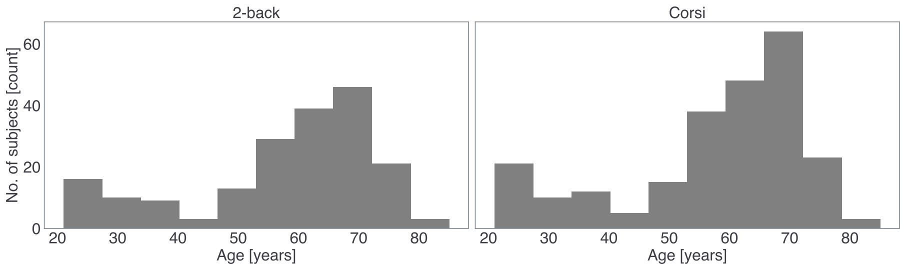


Supplementary Figure 2: The distribution of age of the participants who performed the Corsi block and 2-back tasks.

Supplementary Figure 2: The distribution of age of the participants who performed the Corsi block and 2-back tasks.

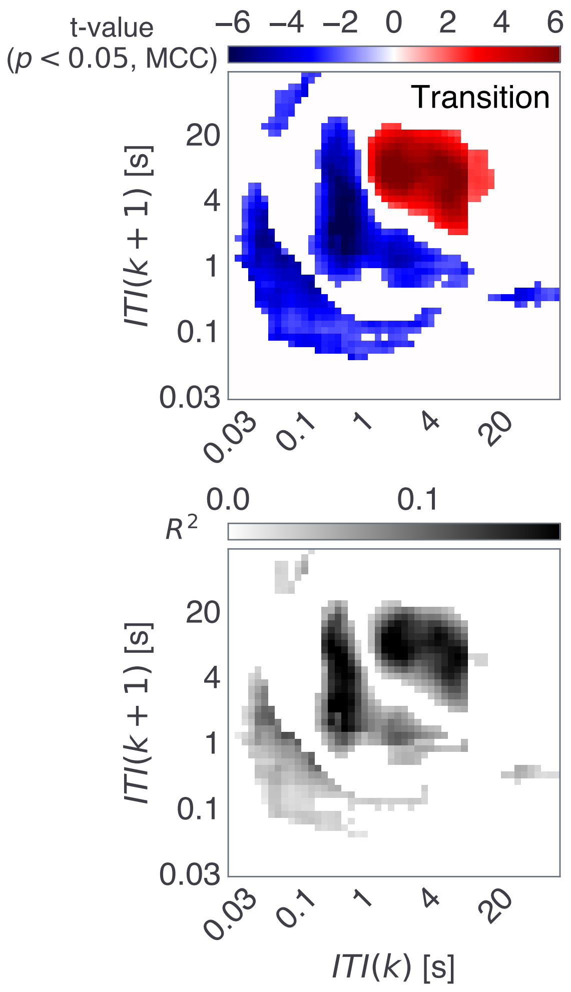


Supplementary Figure 3: Transition JID correlates with the choice reaction time.

Supplementary Figure 3: The t-statistics correspond to the choice reaction time (red-blue image) and the corresponding $R^2$ of the full regression model (incl. gender, gray image). The statistics were corrected for multiple comparisons using two-dimensional clustering.

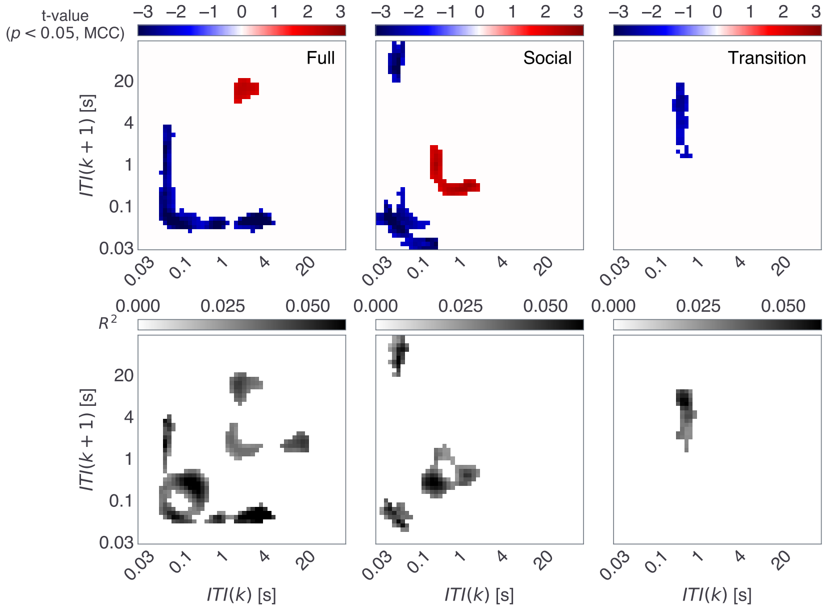


Supplementary Figure 4: Smartphone correlates of simple reaction time.

Supplementary Figure 4: The smartphone behavioral correlates of simple reaction time on the Full, Social, and Transition JIDs. The t-statistics correspond to the cognitive test (red-blue image) and the corresponding $R^2$ of the full regression model (incl. gender, gray image). The statistics were corrected for multiple comparisons using two-dimensional clustering.

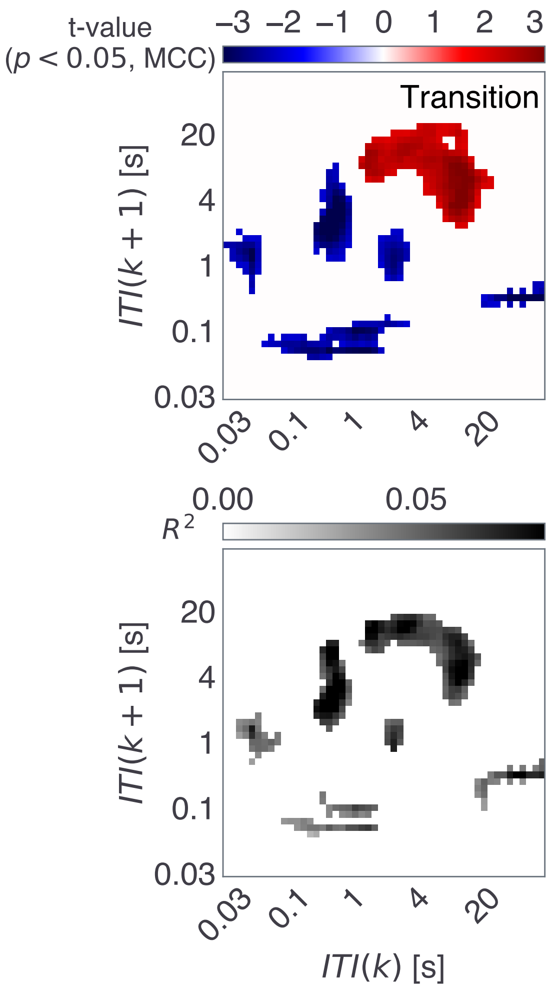


Supplementary Figure 5: Transition JID correlates of global cost.

Supplementary Figure 5: The smartphone behavioral correlates of the global cost on the Transition JID. The t-statistics correspond to the cognitive test (red-blue image) and the corresponding $R^2$ of the full regression model (incl. gender, gray image). The statistics were corrected for multiple comparisons using two-dimensional clustering.

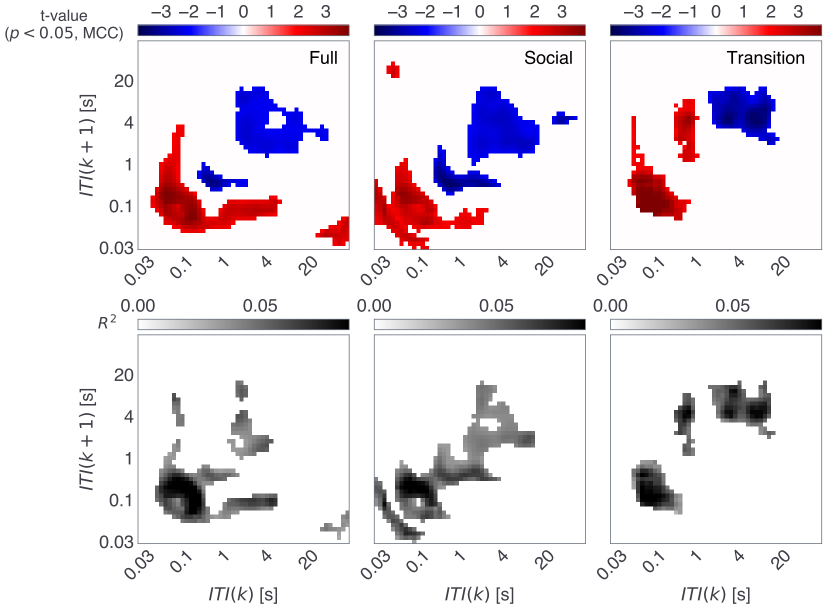


Supplementary Figure 6: Smartphone behavioral correlates of the 2-back test.

Supplementary Figure 6: The smartphone behavioral correlates of 2-back D' on the Full, Social, and Transition JID. The t-statistics correspond to the cognitive test (red-blue image) and the corresponding $R^2$ of the full regression model (incl. gender, gray image). The statistics were corrected for multiple comparisons using two-dimensional clustering.

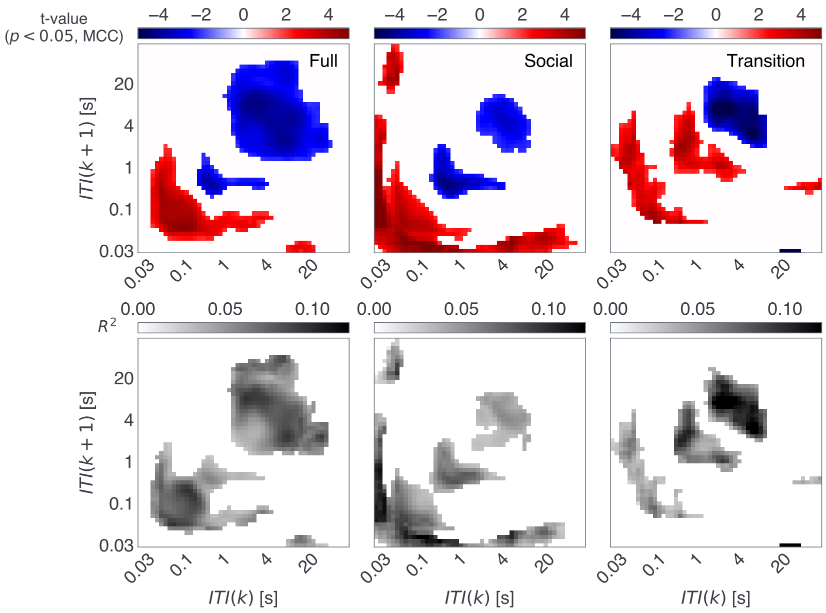


Supplementary Figure 7: Smartphone correlates of Corsi Block.

Supplementary Figure 7: The smartphone behavioral correlates of the span on the Corsi Block task on the Full, Social, and Transition JID. The t-statistics correspond to the cognitive test (red-blue image) and the corresponding $R^2$ of the full regression model (incl. gender, gray image). The statistics were corrected for multiple comparisons using two-dimensional clustering.

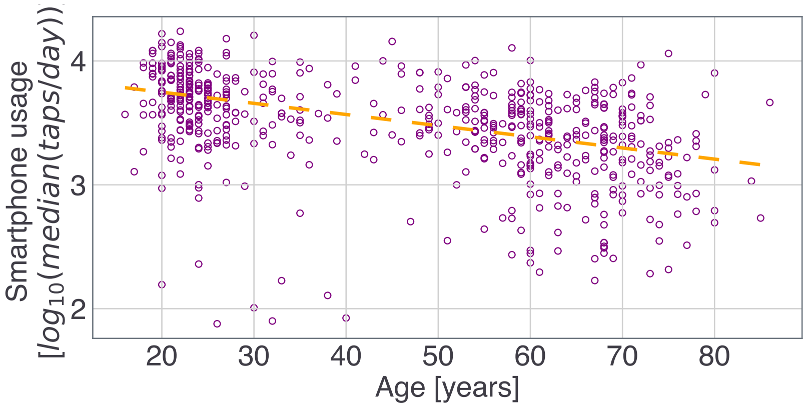


Supplementary Figure 8: Smartphone correlates of age.

Supplementary Figure 8: The amount of smartphone usage correlates with age. Adjusted response plot showing the linear relation between smartphone usage vs. age. Based on robust linear regression against age and gender.

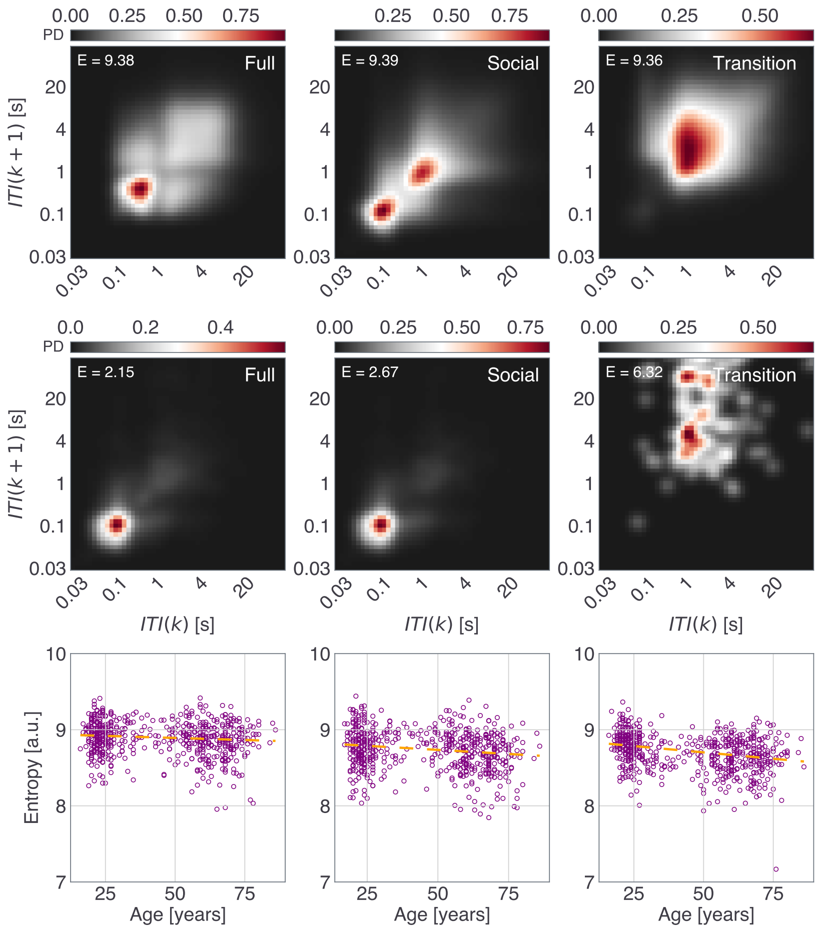


Supplementary Figure 9: Smartphone entropy.

Supplementary Figure 9: The entropy of the JID is related to age. The Full, Social, and Transition JIDs with min and max entropy values in the sampled population. The adjusted response plots show the linear relation between the corresponding entropy vs. age. Based on robust linear regression against age and gender.

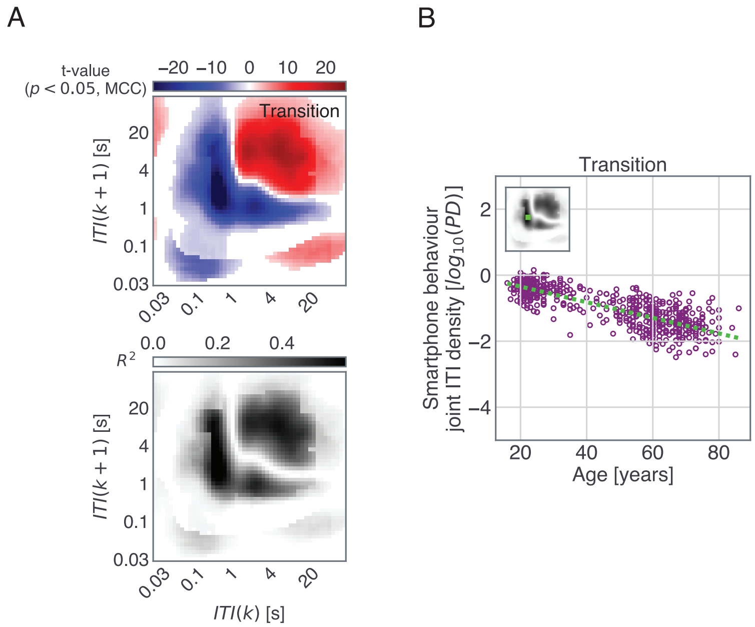


Supplementary Figure 10: Transition JID correlates of age.

Supplementary Figure 10: The correlation between Transition JID and age. Legend same as in Fig. 3 but for Transition JID.

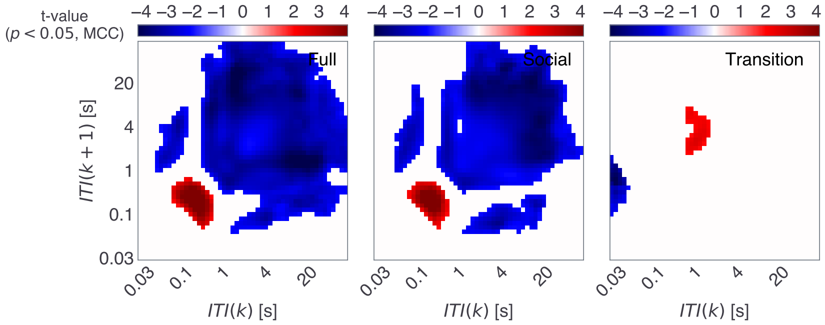


Supplementary Figure 11: Smartphone correlates of gender on the JIDs.

Supplementary Figure 11: The t-statistics (red-blue images) for the variable gender (dummy variable in the regression model including age). Male 1, and female 2 were used as dummy variables. Note, females (2) showed higher probability densities of short consecutive intervals.

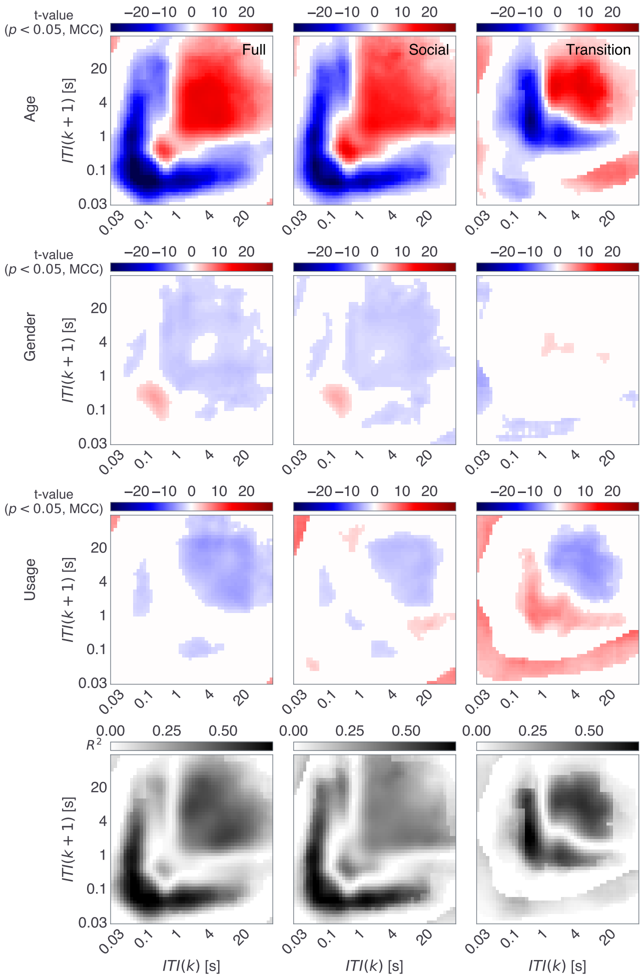


Supplementary Figure 12: A multivariate model including age, gender, and smartphone usage linking to the JIDs.

Supplementary Figure 12: The results of mass univariate regression include the amount of smartphone usage in terms of the median number of interactions per day in addition to age and gender in the mass regressions conducted at each two-dimensional bin. The t-statistics for all of the variables and corresponding $R^2$ of the full model. All statistics were corrected for multiple comparisons using two-dimensional clustering $\alpha = 0.05$.

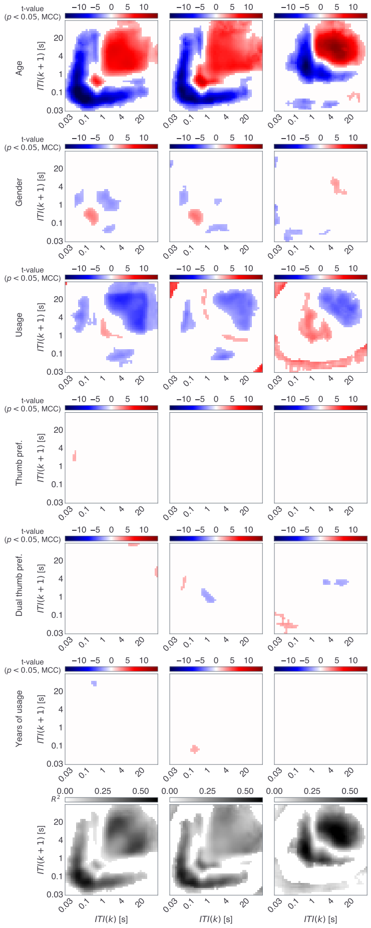


Supplementary Figure 13: Age-related correlates in JIDs remain in the presence of other smartphone use-related variables.

Supplementary Figure 13: We used a multivariate model to explain the inter-individual differences in the JID and this model included age, gender, smartphone usage, thumb preference score, dual thumb preference score, and the years of the smartphone experience. The t-statistics for all of the variables and corresponding $R^2$ of the full model. All statistics were corrected for multiple comparisons using two-dimensional clustering $\alpha = 0.05$.

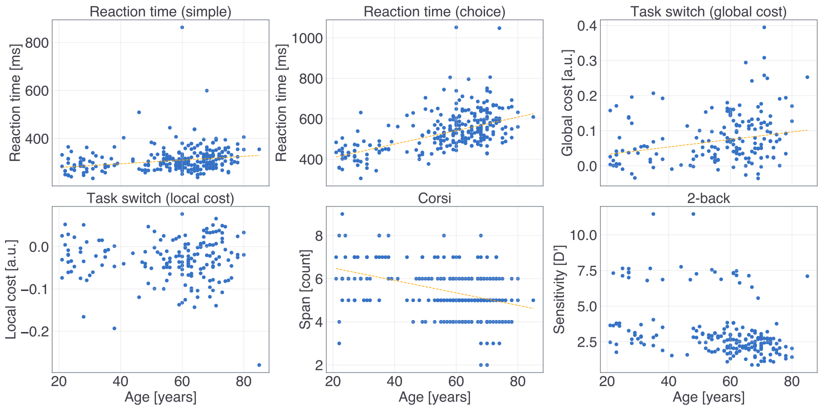


Supplementary Figure 14: Age-related correlates of cognitive tests.

Supplementary Figure 14: The adjusted response plots are based on multivariate regression models containing the variables age and gender. Regression fits with $p < 0.05$ for the variable age are plotted.

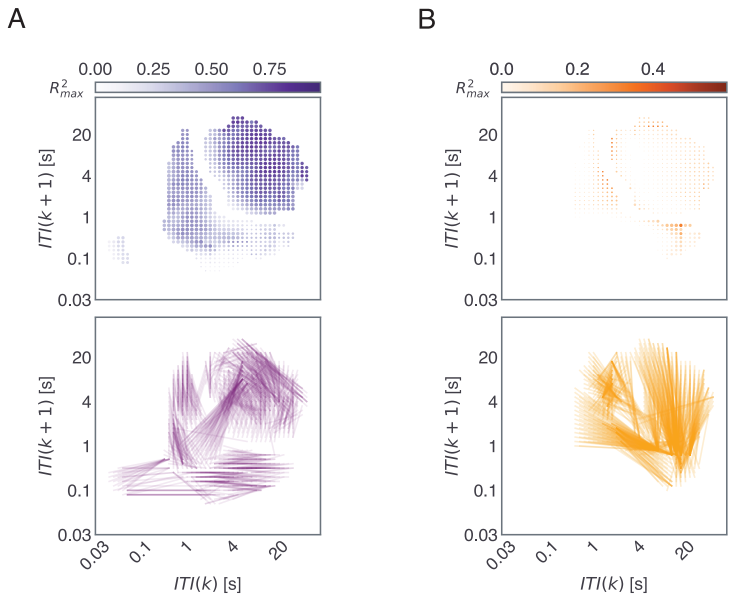


Supplementary Figure 15: Accelerated & decelerated aging captured on Transition JID.

Supplementary Figure 15: (A) Consistent relationships and the corresponding links and (B) inconsistent relationships and the corresponding links. Legend same as in Figure 4 C & D.
